# Engineering a circular riboregulator in *Escherichia coli*

**DOI:** 10.1101/008987

**Authors:** William Rostain, Shensi Shen, Teresa Cordero, Guillermo Rodrigo, Alfonso Jaramillo

## Abstract

Circular RNAs have recently been shown to be important gene expression regulators in mammalian cells. However, their role in prokaryotes remains elusive. Here, we engineered a synthetic riboregulator that self-splice to produce a circular molecule, exploiting group I permuted intron-exon (PIE) sequences. We demonstrated that the resulting circular riboregulator can activate gene expression, showing increased dynamic range compared to the linear form. We characterized the system with a fluorescent reporter and with an antibiotic resistance marker. Thanks to the increased regulatory activity by higher stability, isolation due to self-splicing, and modularity of PIE, we envisage engineered circular riboregulators in further synthetic biology applications.

## Introduction

RNA molecules can regulate gene expression in *trans* through many different mechanisms [1]. The stability of regulatory RNAs, which is difficult to predict, has long been considered as a major bottleneck to achieve efficient activity. This way, two strategies for stabilizing RNA have been identified. First, addition of scaffolds containing binding sites for proteins working as RNA chaperones (e.g., Hfq in bacteria) [2]. This strategy has successfully been used in synthetic small RNAs (sRNAs) to prevent their premature cleavage and degradation by ribonucleases, then improving the activity of the engineered RNA circuits [3, 4]. Second, RNA circularization, which has been shown to enhance RNA stability in living cells [5, 6] by eliminating the exposed 5’ through which degradation by ribonuclease (RNase) E is initiated [7]. To our knowledge, no synthetic circular riboregulator has been engineered to control gene expression in a living cell. Both strategies can nevertheless be found in natural systems. On the one hand, Hfq-associated sRNAs are widespread in enterobacteria [8, 9], and, on the other hand, the development of methods such as CircleSeq [10] has recently shown that circular RNAs are abundant and important regulators in animals [11, 12].

We here propose the engineering of a circular riboregulator in *Escherichia coli*. One way of creating artificially circularized RNAs is to use self-splicing ribozymes [13], which only require GTP and Mg^2+^ for activity. The splicing takes place through two sequential *trans*-esterifications. We here exploited the group I permuted intron-exon (PIE) method [14]. Thanks to permuting the order of the introns and exons and removing the dispensable P6 loop of the bacteriophage T4 *td* ribozyme, the process yields linear introns and circularized exons together with the inserted sRNA (Figure 1a). We designed an RNA sequence so that the resulting structure of the circular RNA (accounting for the exons) does not interfere with the regulatory ability of the molecule.

**Figure 1.**
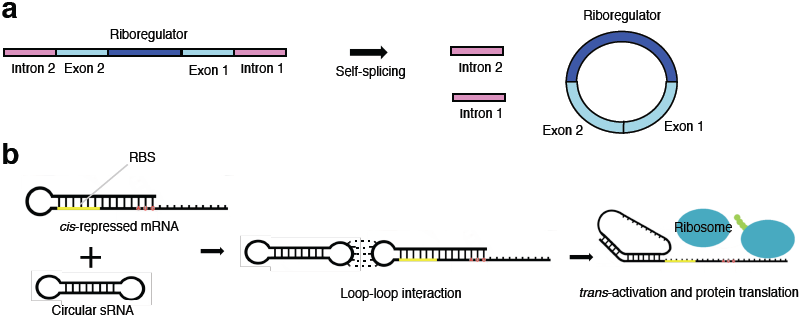
**a**) Diagram of circularization by the PIE method. The riboregulator is inserted between the two exons. **b**) Diagram of riboregulation with a circular sRNA. A *cis*-repressed mRNA is *trans*-activated by the circular sRNA, which induces a conformational change in the 5’ UTR to release the RBS.

Unlike animal regulatory circular RNAs, which have been found to act as sponges with high stability that sequester and inactivate microRNAs (miRNAs), our circular RNA has the capacity to directly interact with a messenger RNA (mRNA) and regulate gene expression. We followed a simple strategy by which the 5’ untranslated region (UTR) of the mRNA *cis*-represses the ribosome-binding site (RBS), and the circular riboregulator interacts with the 5’ UTR to induce a conformational change that releases the RBS and then allows protein translation (Figure 1b) [15].

## Results and Discussion

To design the sequence of circRAJ31, we took a minimal version of our previously engineered sRNA RAJ31 (RAJ31min, Table S1) [16]. This fragment contains the full intermolecular hybridization region with the cognate 5’ UTR. We selected this sRNA because its toehold is within a loop. The predicted secondary structure of circRAJ31 (after splicing and circularization) still exposes the toehold to the solvent.

To ensure the engineered T4-PIE ribozyme retains its splicing ability after insertion of RAJ31min in between the exons, we cloned the engineered circRAJ31 downstream of a T7 promoter and transcribed the RNA *in vitro.* We performed a reverse transcription (RT) followed by a PCR with divergent primers, which would only yield a product provided the RNA molecule is circular, as well as repetitive DNA sequences (Figures 2a,b). The divergent primer-based PCR primary yielded a 112 bp band (size of circRAJ31) and a multiple of 112 bp band in lower extent (Figure 2c), confirming T4-PIE circularization. Sequencing of the RT-PCR product showed the presence of repetitive sequences (Figure 2d), confirming that insertion of RAJ31min within the T4-PIE scaffold yielded a circular RNA *in vitro*, as well as confirming that ligation of the exons happened at the expected site.

**Figure 2.**
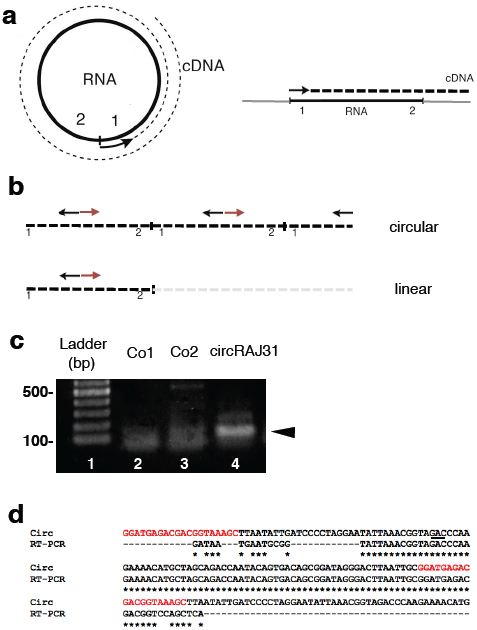
**a**) Diagram of reverse transcription of circularized and linear RNA. **b**) RT-PCR with divergent primers (CircRT-Fw and CircRT-Rv) only yields a product if RNA was circular. **c**) Gel of RT-PCR with primers described above. No RNA template in Control 1 (Co1). No reverse transcriptase in Control 2 (Co2) to check for DNA contamination. Lane 4 shows RTPCR product of circRAJ31. **d**) Sequencing of the RT-PCR product with primer CircRT-Rv. The splicing site is underlined.

We then studied whether circularized riboregulator has an impact in functionality within a cellular environment. For that, we placed circRAJ31 under control of inducible promoter P_LtetO1_ [17] on one plasmid, and the cognate 5’ UTR with the reporter gene under control of inducible promoter P_LlacO1_ [17] on another (Figure 3a). We used a superfolder green fluorescent protein (sfGFP) with a degradation tag (LAA) as reporter. We co-transformed into *E. coli* DH5αZ1 cells, which overexpress the LacI and TetR repressors, providing a tight control of transcription with isopropyl-β-D-thiogalactopyranoside (IPTG) and anhydrotetracycline (aTc). We constructed a non-circularizing mutant (C873U) [18], where a mutation in the P7 loop of the T4 ribozyme (intron 2 in our design) destabilizes the tertiary structure and impinges on circularization [19]. Fluorescence results indicated that T4-PIE circularization allows increasing the dynamic range of a given sRNA (Figure 3b). High gene expression was observed only in the presence of both IPTG and aTc. Whilst the C873U mutant (linear form) gave a fold change of about 2, which is comparable to one reported for the native RAJ31 [16], circRAJ31 was able to reach a fold change of about 6.

**Figure 3.**
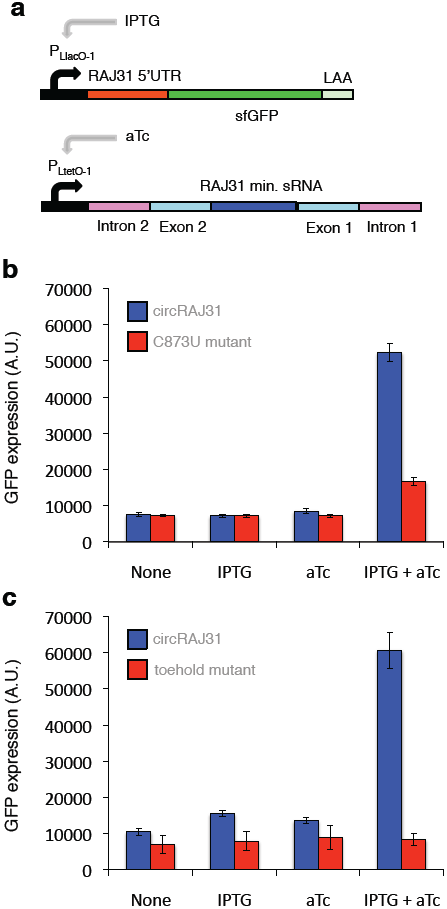
**a**) Scheme of the constructions used: riboregulator circRAJ31 can *trans*-activate a *cis*-repressed gene coding for superfolder green fluorescent protein (sfGFP) with degradation tag (LAA). **b**) Characterization of circRAJ31 and the C873U mutant (in intron 2, to inactivate self-splicing). **c**) Characterization of circRAJ31 and a toehold mutant (in sRNA, to avoid interaction with the mRNA).

To confirm that the exposed toehold in circRAJ31 triggers the activation of the *cis*-repressed sfGFP, we mutated four bases within the toehold of circRAJ31. Toehold mutants are important controls to assess riboregulatory activity and also to derive orthogonal systems [20]. These mutations completely abolished the activity of circRAJ31 (Figure 3c). Importantly, in addition to provide increased regulatory activity by higher stability, T4-PIE circularization may allow the insulation of the sRNA from its genetic context (e.g., to use arbitrary promoters [15]), then increasing the modular design of complex systems with RNA [21].

We then tested the ability of circRAJ31 to tightly control a phenotype. To this end, we used an antibiotic resistance gene (in particular, the *cat* gene, which gives chloramphenicol resistance; Figure 4a). For a tight control, we would need an efficient repression of *cat* expression in absence of the riboregulator together with an efficient activation. We found that circRAJ31 was able to control the phenotype, resulting in growth of colonies on plates containing IPTG and aTc, but not on IPTG or aTc alone (Figure 4b). This suggests that selection for functional sRNAs *in vivo* could be achieved by high-throughput screening.

**Figure 4.**
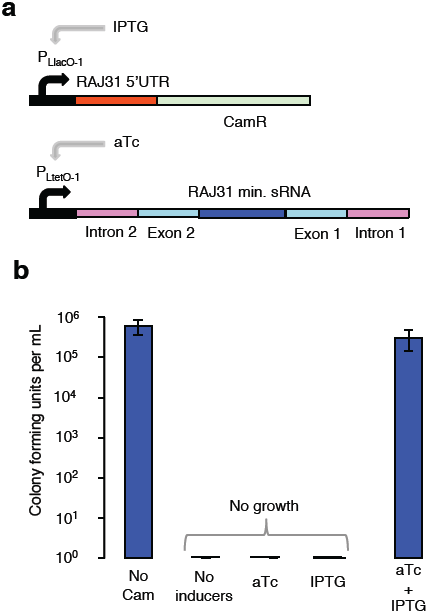
**a**) Scheme of the constructions used: riboregulator circRAJ31 can *trans*-activate a *cis*-repressed gene coding for chloramphenicol acetyltransferase (CamR). **b**) Growth of cells co-transformed with plasmids expressing circRAJ31 and CamR on plates without chloramphenicol (Cam), and on plates with chloramphenicol and different inducers.

To conclude, in this work we investigated the effect of circularization on a synthetic riboregulatory system previously designed by a computational approach (RAJ31) [16]. We encased a minimal version of the sRNA within the T4-PIE scaffold to generate the circularized riboregulator (cirRAJ31). We showed *in vitro* that the system was able to produce a circular sRNA, and also showed that the activation of gene expression *in vivo* (using a fluorescent reporter) was higher than previously reported. Our results suggest that engineered circularization could be a useful strategy to improve the low activity of certain sRNAs, and could potentially be exploited for riboregulation within complex gene networks used for synthetic biology applications.

## Methods

All cloning was performed using *E. coli* DH5α cells (Life Tech.). Riboregulator circRAJ31 (Table S1) was synthesized by IDT in plasmid pIDTSmart, which carries Ampicillin resistance and a pMB1 origin of replication (high copy number). The mRNA (with the 5’ UTR of system RAJ31) was expressed from plasmid pRAJ31m [16], a pSB4K5-derived plasmid, which carries Kanamycin resistance and a mutated pSC101 origin of replication (pSC101m, medium copy number). C873U mutant was constructed using primers tdC873U-Fw and tdC873U-Rv, and toehold mutant with Toe31-Fw and Toe31-Rv (Table S2). DNA sequences were verified using primers VF2 and VR for pRAJ31m and M13-Fw and M13-Rv for pIDTsmart. All chemicals were purchased from Sigma-Aldrich, primers from IDT, and sequencing was done by GATC.

RT-PCR was performed with 2 μL RNA sample, 2 μL primer CircRT-Rv (40 μM) (Table S2), and 4 μL dNTps (10 μM) (final volume 17 μL), with water used instead in controls as necessary, then heated to 65 °C for 5 min, then put on ice. 2 μL tRT x10, 0.5 μL Ribolock, and 0.5 μL M-MuLV-Reverse Transcriptase were then added, before heating to 50 °C for 1 min, then 85 °C for 5 min and finally 20 min at 65 °C. The PCR mix consisted of 4 μL 5x HF, 0.4 μL dNTPs (10 mM), 2μL primer CircRT-Rv (5 μM), 2 μL primer CircRT-Fw (5 μM) (Table S2), 0.6 μL DMSO, 0.2 μL DNA Polymerase Phusion, and 2 μL RT (final volume 20 μL).

For fluorescence assays plasmids were co-transformed in DH5αZ1 cells. Plates and liquid cultures contained Ampicillin, Kanamycin and Spectinomycin to maintain plasmids and Z1 cassette [17]. All M9 medium used was supplemented with 0.8% glycerol, 0.2% casamino acids, 1 μg/mL thiamine, 20 μg/mL uracil and 30 μg/mL leucine and adjusted to pH 7.4. Three single colonies were grown overnight in LB. The next morning, for each condition, 1 mL of M9 medium was inoculated with 5 μL of overnight culture, then grown for 6 h with shaking at 37 °C. 5 μL of refreshed culture were then used to inoculate 195 μL of M9 with antibiotics and appropriate inducers (100 ng/mL aTc, 250 μM IPTG, unless specified otherwise). Three technical replicates were grown for each sample.

Fluorescence measurements were then made in a TECAN Infinite F-500 fluorometer, with 4 cycles/h of shaking with incubation at 37 °C followed by measurement of OD_600_ and fluorescence (excitation 465/35 nm, emission 530/25 nm). For each sample, fluorescence over OD_600_ after background subtraction (corresponding to M9 medium) was taken for the time points closest to OD_600_ = 0.5 [22]. Auto-fluorescence of plain cells was then subtracted. Three technical replicates per three biological replicates (different colonies) were averaged to find a value for each sample, with standard deviation as error bars.

For activation of chloramphenicol resistance assays, three single colonies of co-transformed DH5αZ1 cells were picked, grown overnight in LB. In the morning, they were diluted 1:1000 in LB, grown for 2 h, serially diluted and plated on various inducers. All plates contained 35 μg/mL Kanamycin, 100 μg/mL Ampicillin, 100 μg/mL Spectinomycin as well as 35 μg/mL Chloramphenicol, 100 ng/mL aTc and 250 μM IPTG if appropriate. Plates were then incubated for 20 h at 37 °C and colonies were counted.

## Supporting information

Supporting information is available.

### Notes

The authors declare no competing financial interests.

**Figure S1.**
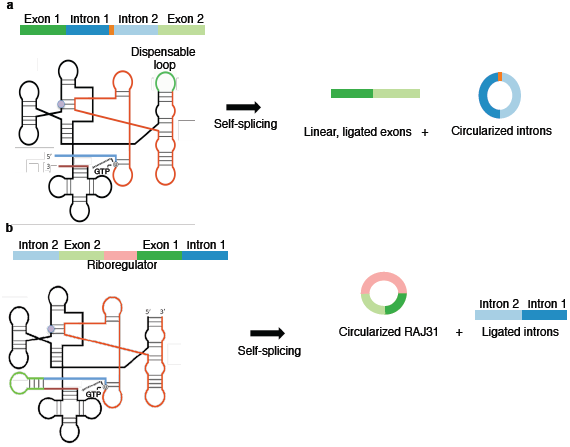
**a**) Diagram and secondary structure of bacteriphage T4 self-splicing ribozyme. Splicing starts with a GTP attack on the Uracil base marked in grey. The C873U mutation (marked in purple) greatly reduces splicing efficiency. **b**) Diagram and structure of PIE ribozyme, created after circular permutation of the original one. The riboregulator is inserted between the two exons, and the sequence starts in the truncated P6 stem. Splicing of PIE ribozymes produces two linear RNAs composed of the introns and a circularized riboregulator.

**Figure S2.**
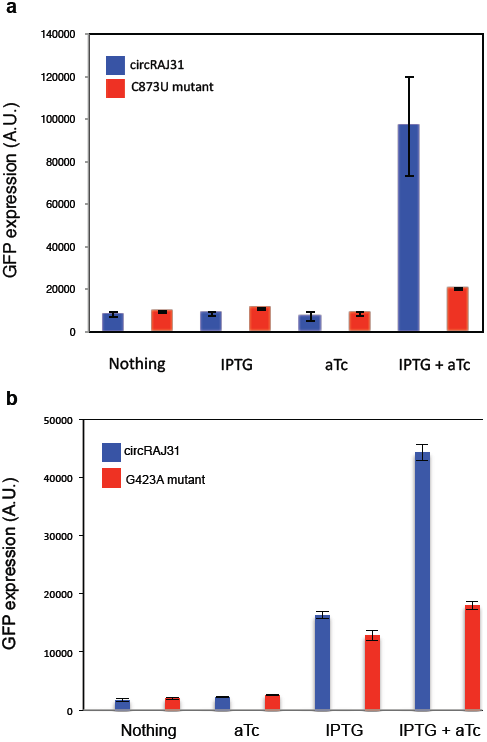
Characterization of circRAJ31 and **a**) the C873U mutant (IPTG concentration = 0.25 mM) and b) G423A mutant (IPTG concentration = 1mM). In both cases, the reporter is a superfolder green fluorescent protein (sfGFP) without degradation tag. In this case, the time points for analysis were closest to OD_600_ = 0.45 and auto-fluorescence of plain cells was not subtracted.

**Figure S3.**
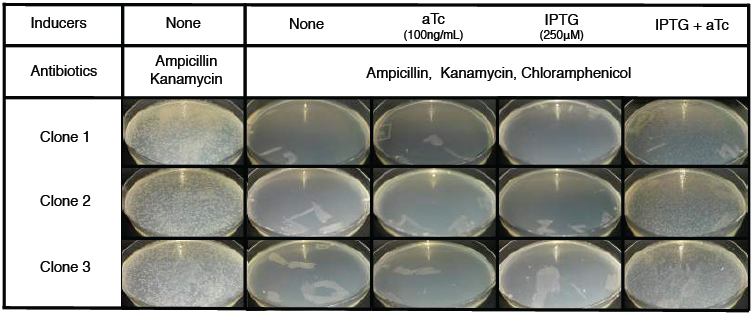
Growth of cells co-transformed with plasmids expressing circRAJ31 and CamR on plates without chloramphenicol (Cam), and on plates with chloramphenicol and different inducers.

**Table S1.**
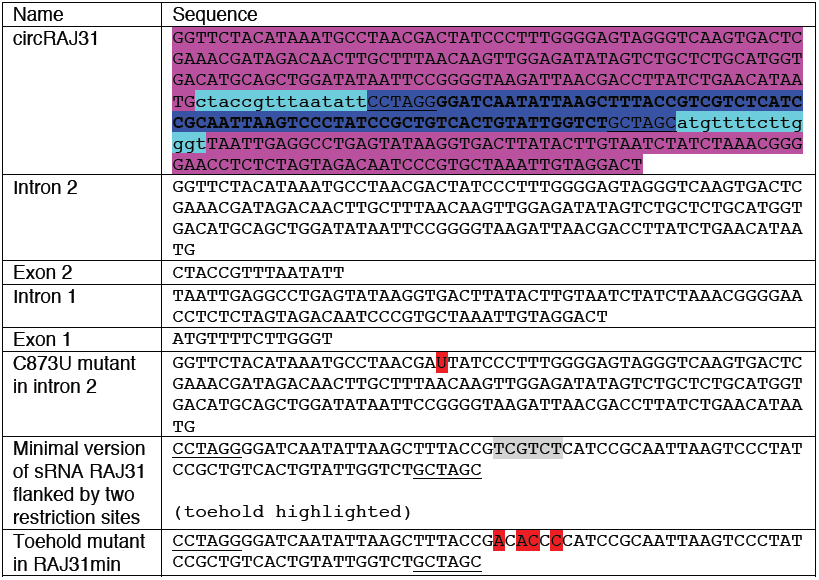
Sequences of the system. To express circRAJ31, we used the P_LtetO1_ promoter and the *rrnC* terminator.

**Table S2.**
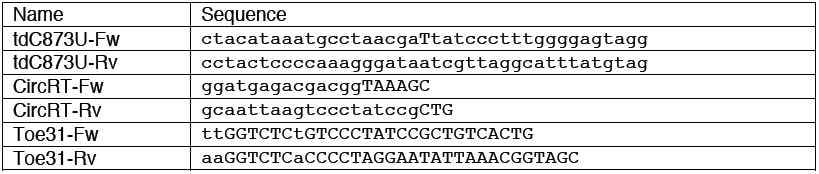
Sequences of the primers used in this work.

## Acknowledgment

Work funded by the grants FP7-ICT-610730 (EVOPROG) and PROMYS (FP7-KBBE-613745) to AJ. WR is supported by a DGA graduate fellowship, and GR by the AXA Research Fund. We thank D. Christiany for his help in the lab. We greatly thank S. Umekage for providing the PIE sequence, and for very helpful advice and discussions.

## References

1. Qi, L. S., and Arkin, A. P. (2014) A versatile framework for microbial engineering using synthetic non-coding RNAs, Nat. Rev. Microbiol. 12, 341–354.

2. Moll, I., Afonyushkin, T., Vytvytska, O., and Kaberdin, V. R. (2003) Coincident Hfq binding and RNase E cleavage sites on mRNA and small regulatory RNAs, RNA 9, 1308–1314.

3. Na, D., Yoo, S. M., Chung, H., Park, H., Park, J. H., and Lee, S. Y. (2013) Metabolic engineering of Escherichia coli using synthetic small regulatory RNAs. Nat. Biotechnol. 31, 170–174.

4. Sakai, Y., Abe, K., Nakashima, S., Yoshida, W., Ferri, S., Sode, K., and Ikebukuro, K. (2013) Improving the gene-regulation ability of small RNAs by scaffold engineering in Escherichia coli, ACS Synth. Biol. 3, 152–162.

5. Umekage, S., and Kikuchi, Y. (2009) In vitro and in vivo production and purification of circular RNA aptamer, J. Biotechnol. 139, 265–272.

6. Umekage, S., Uehara, T., Fujita, Y., Suzuki, H. & Kikuchi, Y. (2012) In Vivo Circular RNA Expression by the Permuted Intron-Exon Method. Innov. Biotechnol. 75–90.

7. Mackie, G. A. (1998) Ribonuclease E is a 5’-end-dependent endonuclease, Nature 395, 720–723.

8. Zhang, A., Wassarman, K. M., Rosenow, C., Tjaden, B. C., Storz, G., and Gottesman, S. (2003) Global analysis of small RNA and mRNA targets of Hfq, Mol. Microbiol. 50, 1111–1124.

9. Sittka, A., Lucchini, S., Papenfort, K., Sharma, C. M., Rolle, K., Binnewies, T. T., Hinton, J. C. D., and Vogel, J. (2008) Deep sequencing analysis of small noncoding RNA and mRNA targets of the global post-transcriptional regulator, Hfq, PLoS Genet. 4, e1000163.

10. Jeck, W. R., and Sharpless, N. E. (2014) Detecting and characterizing circular RNAs, Nat. Biotechnol. 32, 453–461.

11. Hansen, T. B., Jensen, T. I., Clausen, B. H., Bramsen, J. B., Finsen, B., Damgaard, C. K., and Kjems, J. (2013) Natural RNA circles function as efficient microRNA sponges, Nature 495, 384–388.

12. Memczak, S., Jens, M., Elefsinioti, A., Torti, F., Krueger, J., Rybak, A., Maier, L., Mackowiak, S. D., Gregersen, L. H., Munschauer, M., Loewer, A., Ziebold, U., Landthaler, M., Kocks, C., le Noble, F., and Rajewsky, N. (2013) Circular RNAs are a large class of animal RNAs with regulatory potency, Nature 495, 333–338.

13. Cech, T. R. (1990) Self-Splicing of Group I Introns, Annu. Rev. Biochem. 59, 543–568.

14. Ford, E., and Ares, M. (1994) Synthesis of circular RNA in bacteria and yeast using RNA cyclase ribozymes derived from a group I intron of phage T4, Proc. Natl. Acad. Sci. USA 91, 3117–3121.

15. Isaacs, F. J., Dwyer, D. J., Ding, C., Pervouchine, D. D., Cantor, C. R., and Collins, J. J. (2004) Engineered riboregulators enable post-transcriptional control of gene expression, Nat. Biotechnol. 22, 841–847.

16. Rodrigo, G., Landrain, T. E., and Jaramillo, A. (2012) De novo automated design of small RNA circuits for engineering synthetic riboregulation in living cells, Proc. Natl. Acad. Sci. USA 109, 15271–15276.

17. Lutz, R., and Bujard, H. (1997) Independent and tight regulation of transcriptional units in Escherichia coli via the LacR/O, the TetR/O and AraC/I1-I2 regulatory elements, Nucleic Acids Res. 25, 1203–1210.

18. Belfort, M., Chandry, P. S., and Pedersen-Lane, J. (1987) Genetic delineation of functional components of the group I intron in the phage T4 td gene, Cold Spring Harb. Symp. Quant. Biol. 52, 181–192.

19. Brion, P., Michel, F., Schroeder, R., and Westhof, E. (1999) Analysis of the cooperative thermal unfolding of the td intron of bacteriophage T4, Nucleic Acids Res. 27, 2494–2502.

20. Rodrigo, G., Landrain, T. E., Majer, E., Daròs, J.-A., and Jaramillo, A. (2013) Full design automation of multi-state RNA devices to program gene expression using energy-based optimization, PLoS Comput. Biol. 9, e1003172.

21. Lou, C., Stanton, B., Chen, Y. J., Munsky, B., and Voigt, C. A. (2012) Ribozyme-based insulator parts buffer synthetic circuits from genetic context, Nat. Biotechnol. 30, 1137–1142.

22. Wang, B., Kitney, R. I., Joly, N., and Buck, M. (2011) Engineering modular and orthogonal genetic logic gates for robust digital-like synthetic biology, Nat. Commun. 2, 508.

